# Targeting DNA Methylation Reactivates Type I Interferon Signalling in Bone-Metastatic Breast Cancer

**DOI:** 10.64898/2026.06.28.735052

**Authors:** Joan So, Thomas B Chadwick, Miriam Fuentes-Guirado, Laura Vojtech, Lap Hing Chi, Braydon Meyer, Ruth Pidsley, Nicole Haynes, Hugh Chalmers, Aliye Tabatabaee, Amina Ismail, Karla J. Cowley, Kaylene J. Simpson, Clare Stirzaker, Belinda S. Parker

**Affiliations:** Sir Peter MacCallum Department of Oncology, The University of Melbourne, Parkville, Victoria, Australia; Cancer Evolution and Metastasis Program, Peter MacCallum Cancer Centre, Melbourne, Victoria, Australia; Epigenetics Research Laboratory, Cancer Ecosystems Program, Garvan Institute of Medical Research, Sydney, NSW, 2010, Australia; School of Clinical Medicine, UNSW Medicine and Health, Sydney, NSW, 2010, Australia; Victorian Centre for Functional Genomics, Peter MacCallum Cancer Centre, Parkville, VIC, Australia

## Abstract

Bone metastasis remains a major clinical challenge in advanced breast cancer. Downregulation of type I interferon signalling, a critical immunomodulatory pathway in anti-cancer immunity and disease progression, is a defining feature of this process. Here, we utilised an IFN-reporter system to perform unbiased epigenetic compound screens to identify agents that could restore type I IFN signalling. This screen identified Decitabine, a DNA hypomethylating agent, that enhanced tumor immunogenicity across a broad range of mouse and human breast cancer cell lines, including bone-derived lines. Mechanistically, suppression of interferon-stimulated genes is highly correlated with elevated DNMT1 expression in bone metastasis compared with primary tumor, in both mouse models and matched human samples. Decitabine treatment was sufficient to reactivate interferon stimulated genes in bone-derived 4T1.2 cell lines via hypomethylation of type I interferon pathway gene promoter regions. Correspondingly, in the syngeneic 4T1.2 metastasis mouse model, Decitabine treatment conferred a survival benefit and reduced metastatic potential, particularly in bone. Our findings reveal DNA methylation as a key regulator of the transcriptional programs underlying bone metastasis, providing mechanistic insight into how Decitabine reactivates type I interferon signalling and reduces metastatic potential, highlighting epigenetic reprogramming as a promising approach for targeting metastatic breast cancer.

**STATEMENT OF SIGNIFICANCE:** Bone metastasis remains a major clinical challenge and a key mechanism of progression to bone is the suppression of tumor-inherent type I interferon signalling. We identified Decitabine, a DNA hypomethylating agent, as a promising therapeutic agent to enhance tumor immunogenicity across a broad range of breast cancer cell lines, including bone metastasis-derived lines. Our findings support DNA methylation as a key regulator of transcriptional programs associated with bone metastatic progression, and provide mechanistic insight into how Decitabine reactivates type I interferon signalling and reduces metastatic potential *in vivo*. These results highlight epigenetic reprogramming as a promising approach for targeting metastatic breast cancer.

## INTRODUCTION

Breast cancer (BCa) is the most commonly diagnosed cancer and leading cause of cancer-related death in females globally^1^. Recent years have shown vast improvements in the detection and treatment of early-stage BCa, from national screening programs to the integration of novel therapies such as immune checkpoint inhibition^1,2^ and HER-targeted agents, including next generation monoclonal antibodies and antibody drug conjugates with markedly improved efficacy^3,4^. Despite this, across subtypes approximately 20-30% of patients progress to stage IV and develop metastatic disease, for which there are limited curative treatment options^5^. Bone metastasis is the most common site of spread, and due to the rapid resorption and destruction of bone patients are at risk of skeletal-related events (SREs), including reduced mobility, pathological fractures, hypercalcemia and spinal-cord fractures; reducing quality of life and increasing both morbidity and mortality^6–9^. With the global healthcare burden that metastatic BCa entails, novel methods of treating and curing bone metastasis are desperately needed.

During the process of metastasis, tumor cells undergo a number of changes to adapt and survive the metastatic cascade; involving epithelial-to-mesenchymal (EMT) transition to facilitate extravasation into the blood stream, survival from immune elimination, and intravasation and colonization into distal organs^10^. Notably, at the time of primary tumor diagnosis, it is estimated that around 40% of BCa patients already harbour micro-metastases within the bone marrow^11–13^, and so the understanding and treatment of metastatic cells may be important even at an early-stage diagnosis. A key feature that differentiates bone metastatic BCa cells from their primary tumor counterpart is the loss of tumor intrinsic type I interferon (IFN) signalling: a critical immunomodulatory pathway well characterised in anti-cancer immunity and disease progression^10,14,15^. Our laboratory has previously characterised the loss of transcription factors IFN regulatory factors 7 and 9 (IRF7 and IRF9) and associated IFN stimulated gene (ISG) signatures in breast and prostate cancer bone metastasis across both mouse models and patient derived samples^16–19^. We also demonstrated that loss of the Interferon-alpha/beta Receptor Subunit 1 (IFNAR1) was sufficient to drive bone metastasis in 4T1 and 66cl4 murine models of BCa^20^. Furthermore, loss of IFN and IRF7 associated signatures in breast primary tumors alone predicted the risk of distal metastasis to the bone in patients^16,21^. The expression of type I IFN signalling is critical for response to chemotherapy^22^: where the expression of IRF9 can predict long-term neoadjuvant chemotherapeutic response in TNBC^17^ and overexpression of IRF7 and associated ISGs has been shown to sensitise tumors to both DNA-damaging chemotherapy and anti-PD1 checkpoint inhibition, prolonging metastasis-free survival in mouse models of BCa^16,22,23^. Despite the importance of this pathway, IFN-based therapies have previously failed (reviewed in ^14,15^) primarily due to toxicities associated with chronic and systemic IFN expression and a lack of understanding as to what drives the metastasis-specific repression.

While genomic and oncogenic alterations can drive the loss of type I IFN signalling in cancer, tumor cells frequently suppress the pathway by epigenetically silencing key components: from IFN/IRF genes^24,25^, pattern recognition receptors such as cyclic GMP−AMP synthase (cGAS) and stimulator of IFN genes (STING)^26–28^, and downstream ISGs^29,30^. In models of BCa, several mechanisms of epigenetic IFN suppression have been described, including upregulation of DNA methylation through DNA (cytosine-5)-methyltransferase 1 (DNMT1) and DNMT3A/B^31,32^, RNA methyltransferases such as methyltransferase-like 3 (METTL3)^33^, histone methyltransferases like SET domain bifurcated histone lysine methyltransferase 1 (SETDB1)^34^ and histone deacetylases 1-3 (HDAC1-3) all being implicated in facilitating immune evasion^35–37^ (reviewed in ^15^). Genetic or pharmacological ablation of these regulators can restore IFN signalling through the derepression of associated genes directly, and therefore may restore resistance to therapeutics that rely on this signalling, including DNA damaging agents. Furthermore, transcription of normal silenced retroelements through DNMT inhibitors or other agents may elicit an IFN response through the generation of cytosolic dsRNA, in a phenomenon known as “viral mimicry”^38,39^. However, while an increasing number of epigenetic agents are being tested in the primary breast tumor setting^40^, little is known regarding the role that these suppression pathways play in the characteristic IFN loss and treatment resistance observed in bone metastasis, and if they can be targeted therapeutically.

In the present study, we utilised a high-throughput epigenetic drug screen on multiple primary tumor and bone-metastasis derived BCa cell lines to identify novel inducers of IFN signalling. Using an IFN stimulated response element (ISRE) reporter as the primary readout, we found the DNMT inhibitor and hypomethylating agent Decitabine (DAC) as a potent inducer of ISRE activity across human and mouse BCa lines, and an effective anti-metastatic agent *in vivo*. Furthermore, we characterised the distinct transcriptional profile and DNA methylation landscape in bone metastasis that drives its immune suppression. Our results suggest that DNA methylation is a key driver of IFN suppression in bone metastasis and that targeting this mechanism could be an effective strategy for late stage BCa.

## RESULTS

### Pharmacological Screening Identifies Context-Dependent Inducers of Interferon Signalling in BCa Cells

To identify candidate mechanisms regulating repression of type I IFN signatures in BCa cells, we performed a drug screen across a panel of mouse and human cell lines with distinct metastatic potential. These included highly metastatic mouse lines 4T1.2_PAR (parental 4T1.2 cell lines), 4T1.2_PT (derived from primary tumors), and 4T1.2_BONE (derived from bone metastases), as well as the moderately metastatic J110 line. Human models comprised highly metastatic MDA-MB-231, moderately metastatic CAL-120, and low metastatic localised T47D. Cells were transduced with an ISRE reporter to monitor IFN activity, and reporter induction was systematically assessed 72 hours following drug treatment (Fig. 1A).

**Figure 1.**
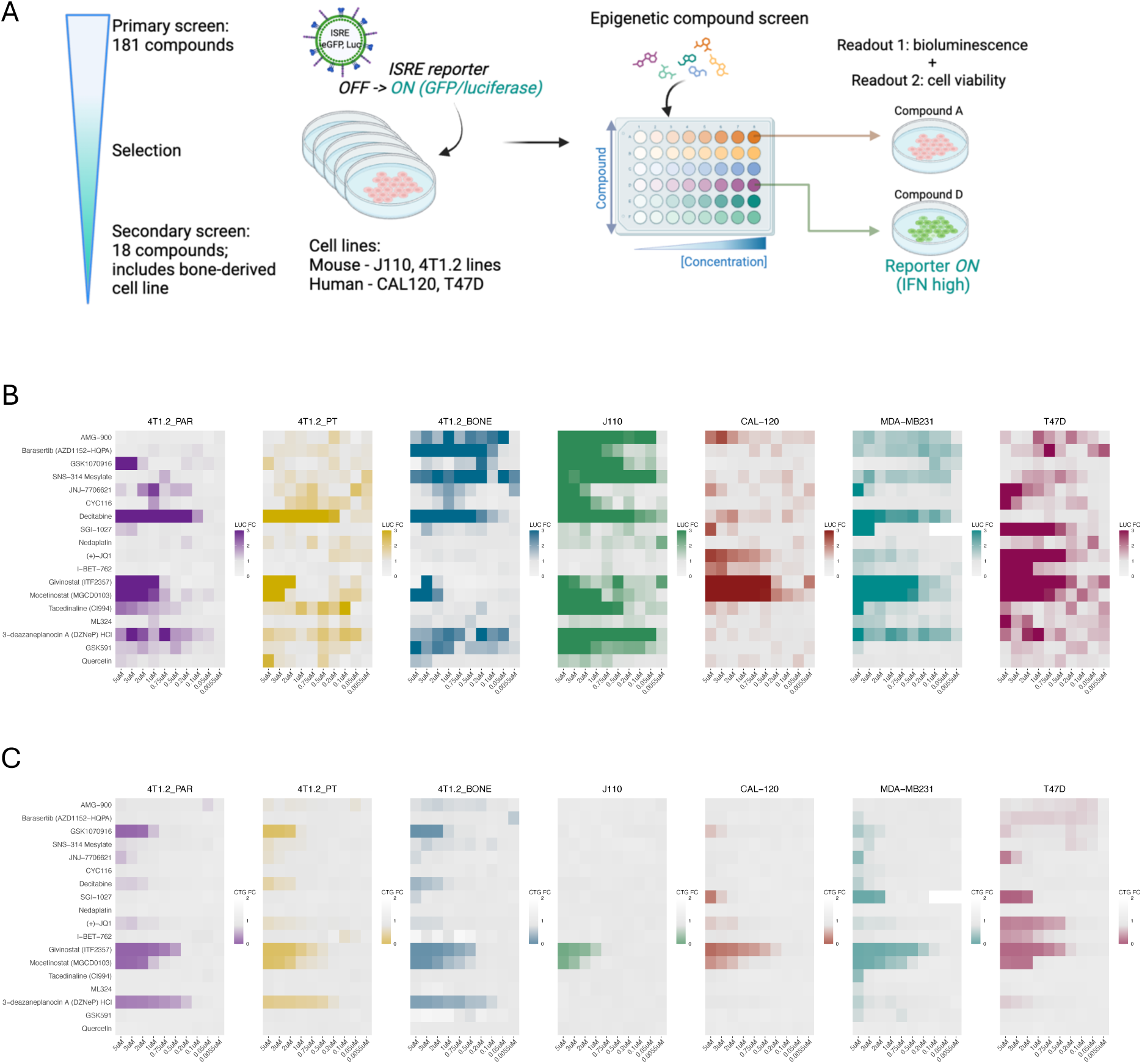
Pharmacological screening identifies context-dependent inducers of IFN signalling in BCa models. **(A)** Experimental workflow of the ISRE reporter-based drug screen performed across mouse and human BCa cell lines with varying metastatic potential. **(B)** Heatmap showing ISRE reporter activity following treatment with the indicated compounds for 72 hours. Values represent fold-change relative to vehicle-treated controls. DAC, Mocetinostat and DZNep-HCl emerged as the strongest activators of IFN signalling across multiple models. **(C)** Corresponding heatmap showing cell viability following treatment with the indicated compounds for 72 hours. Values are expressed relative to vehicle-treated controls. DAC induced ISRE activity with limited cytotoxicity, whereas Mocetinostat and DZNep-HCl reduced cell viability.

Top candidates from the full screen of 181 compounds were taken through to a validation screen (Supp. Fig. 1A-B), which narrowed the pool down to those with the most activity. In mouse cell lines, DAC, Mocetinostat, and DZNep-HCl consistently induced ISRE reporter activity in a dose-dependent manner (Fig. 1A), with DAC as the only compound that specifically enhanced IFN signalling at lower doses without impacting cell viability (Fig. 1B-C). Notably, 4T1.2_BONE cells exhibited increased sensitivity to the Aurora kinase inhibitors Barasertib and SNS-314 mesylate compared to matched 4T1.2_PT cells with minimal impact on cell viability, indicating a bone-specific response. In human cell lines, Mocetinostat and DZNep-HCl similarly induced ISRE activity and cell death in a dose-dependent manner across models. In contrast, DAC elicited robust ISRE activation in highly metastatic MDA-MB-231 cells but weaker responses in less metastatic CAL-120 and T47D cells, suggesting preferential activity in metastatic contexts. Barasertib and SNS-314 Mesylate induced modest ISRE activation in MDA-MB-231, with variable and inconsistent effects observed in CAL-120 and T47D. Collectively, these findings identify DAC as a potent inducer of ISRE activity across multiple models, highlighting its therapeutic potential to reactivate IFN signalling, particularly in metastatic disease settings.

### DAC induces ISG mRNA expression as a single agent and enhances ISRE response to Poly (I:C)

To validate enhanced transcription of canonical ISGs themselves, we examined their expression in matched PT- and bone-metastatic 4T1.2 cell lines, before and after DAC treatment (Fig. 2A). In 4T1.2_BONE compared to 4T1.2_PT, canonical ISG mRNA were downregulated (Fig 2B), including an 85.8% reduction in *Irf7* (p < 0.0001), 39.9% reduction in *Irf9* (p = 0.0036), 72.9% reduction in *Oas1* (p < 0.0001) and a 54.2% reduction in *Psmb9* (p = 0.0002). *Tap1* was unchanged between groups (p = 0.4565). Treatment of 4T1.2_PT and 4T1.2_BONE cell lines with 250nM DAC or DMSO for 72 hours resulted in enhanced expression of ISGs, an effect that was more pronounced in the lines derived from bone. DAC treatment increased the expression of *Psmb9* and *Tap1* in both the 4T1.2_PT and BONE lines (Fig. 2F-G and 2K-L), however the 4T1.2_BONE line also demonstrated increases in *Irf9* and greater trends in *Irf7* and *Oas1* (Fig. 2H-J).

**Figure 2.**
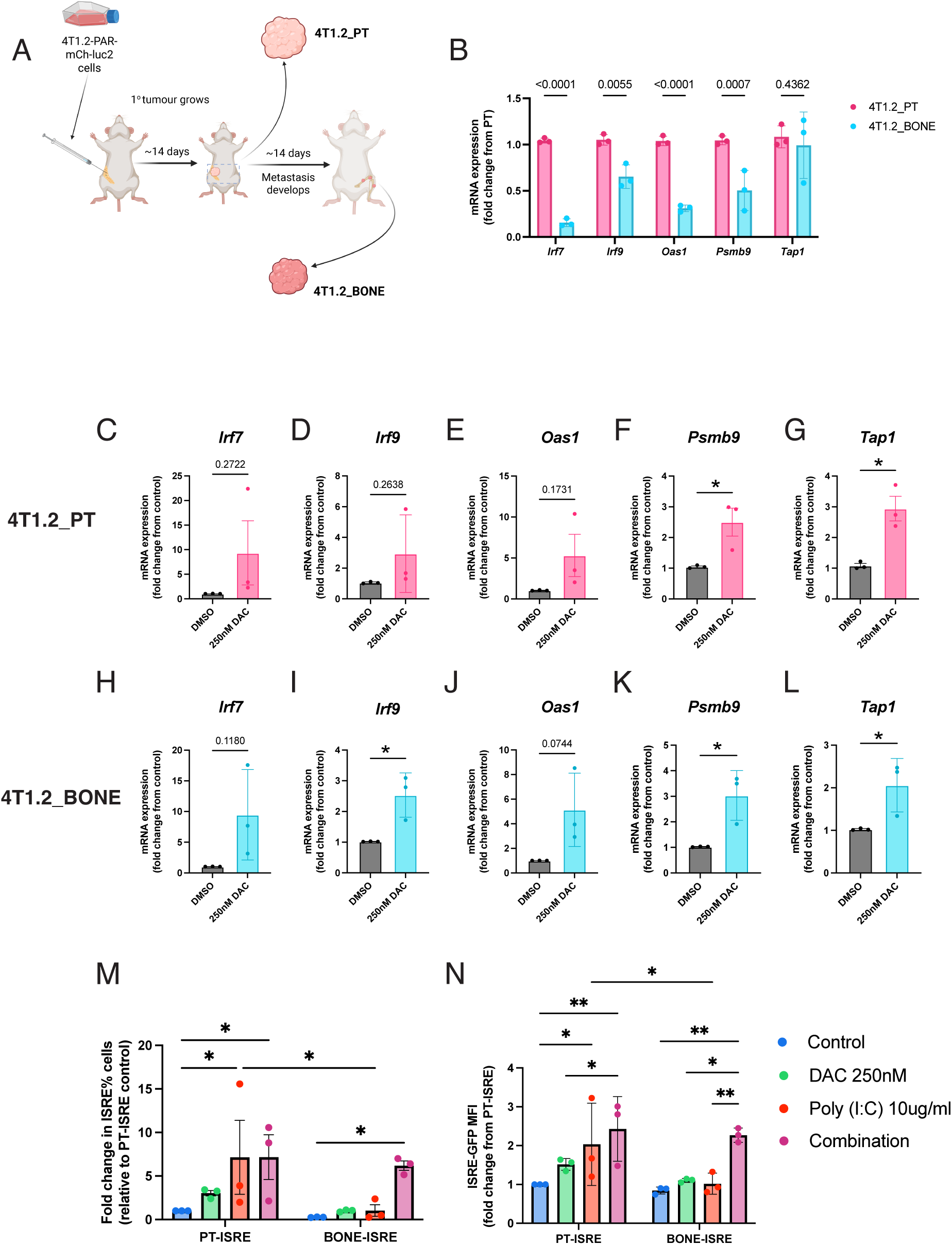
Decitabine induces ISG expression in both primary tumor and bone metastasis-derived 4T1.2 cells and enhances bone metastasis response to Poly (I:C) **(A)** The parental 4T1.2 cells were injected into mammary fat pad of syngeneic Balb/c mice to initiate primary tumor development and spontaneous bone metastasis. Primary tumor was resected on day 14 post-injection (∼300mm^3^) to establish the 4T1.2_PT cell line. Cancer metastases were extracted from the femur on day 28 post-injection to establish the 4T1.2_BONE cell line. Both lines were cultured *in vitro* and underwent two passages before experimental studies were undertaken. **(B)** qRT-PCR analysis of canonical ISG mRNA expression between 4T1.2_PT and 4T1.2_BONE cell lines, revealing widespread ISG downregulation in the BONE compared to PT. **(C-L)** After 72 hours of 250 nM DAC therapy, canonical ISG mRNA abundance was analysed between DMSO and DAC groups, measuring *Irf7, Irf9, Oas1, Psmb9* and *Tap1* expression in 4T1.2_PT (**C-G)** and 4T1.2_BONE cell lines **(H-L)**. **(M-N)** The pGF-ISRE-GFP-luc2 reporter expression was determined by relative fold change of percentage of positive cells **(M)** and median fluorescence intensity (MFI) **(N)** quantified at 72-hour post-DAC therapy (dose) and/or 24-hour post Poly (I:C) treatment (10μg/ml) by flow cytometry measuring ISRE-GFP expression to determine synergistic effects of DAC with an IFN inducer. Data represented as mean +/- SEM. Statistical analysis was performed using a repeated-measures two-way ANOVA with Tukey’s post-hoc test for **(B, M, N)** and a two-tailed, unpaired, parametric t-test for (**C-L).** P-values was calculated using two-tailed, unpaired, non-parametric t-test. * P < 0.05, **P < 0.01. n=3 for all, biological replicates were determined but individual freeze-thaws and individual analysis run for both qRT-PCR and flow cytometry. ISG, interferon-stimulated gene; DAC: decitabine; IFN, interferon.

Furthermore, to test DAC’s impact on ISRE-reporter activity in combination with an IFN-inducer, we treated 4T1.2_PT-ISRE and 4T1.2_BONE-ISRE cells with either 72-hour DAC or 24-hour Poly (I:C) (a TLR3 agonist) or a combination (Fig 2M-N). In the PT-ISRE line, DAC did not ISRE positive cell proportions or MFI. As expected for the primary tumor line, the addition of Poly I:C increased both markers, however addition of decitabine in combination did not enhance this. Conversely, the bone metastatic lines relied on the combination for restored ISRE reporter signaling, whereby DAC and Poly (I:C) alone had no impact, yet the DAC and Poly (I:C) combination resulted in over a 2-fold increase. Furthermore, there was over a 6-fold decrease in percentage ISRE positivity and MFI post Poly (I:C) in BONE-ISRE groups compared to the PT-ISRE lines. Studies incorporating additional cell lines and technical replicates are ongoing and will be reported in the final manuscript.

### DAC Reactivates Interferon Signalling in Bone- and Primary-Derived Tumor Cells with Distinct Transcriptional Programs

To characterise transcriptional changes associated with bone colonisation, we performed bulk RNA sequencing using 4T1.2 cell lines isolated from bone metastases (Bone) and matched primary mammary fat pad tumors (PT). To determine whether tissue-derived lines retained stable transcriptional divergence from the parental population, we also sequenced the parental 4T1.2 cell line (Parental). Multidimensional scaling analysis demonstrated clear separation among Parental, PT, and Bone cell populations, confirming stable transcriptional reprogramming following tissue adaptation despite passaging *in vitro*.

Comparative analysis revealed 844 genes significantly downregulated in Bone relative to PT, including multiple canonical ISGs, such as *Irf7, Usp18, Zbp1, Oas1a, Oas1g,* and *Mx2* (Fig. 3A-B), consistent with the targeted gene analysis in Fig 2B and our previous observations^16^. In contrast, 717 genes were significantly upregulated in Bone cells, including core DNA methylation maintenance machinery (*Dnmt1* and *Uhrf1*) and proliferation-associated regulators (*Pcna* and *Pclaf*) (Fig 3A), in line with prior clinical reports linking DNMT1 overexpression with metastatic progression. In agreement with the individual gene expression, pathway enrichment analysis demonstrated strong suppression of interferon-related pathways, alongside coordinated activation of MYC and E2F proliferative programmes in Bone cells compared with PT (Fig. 3C).

**Figure 3.**
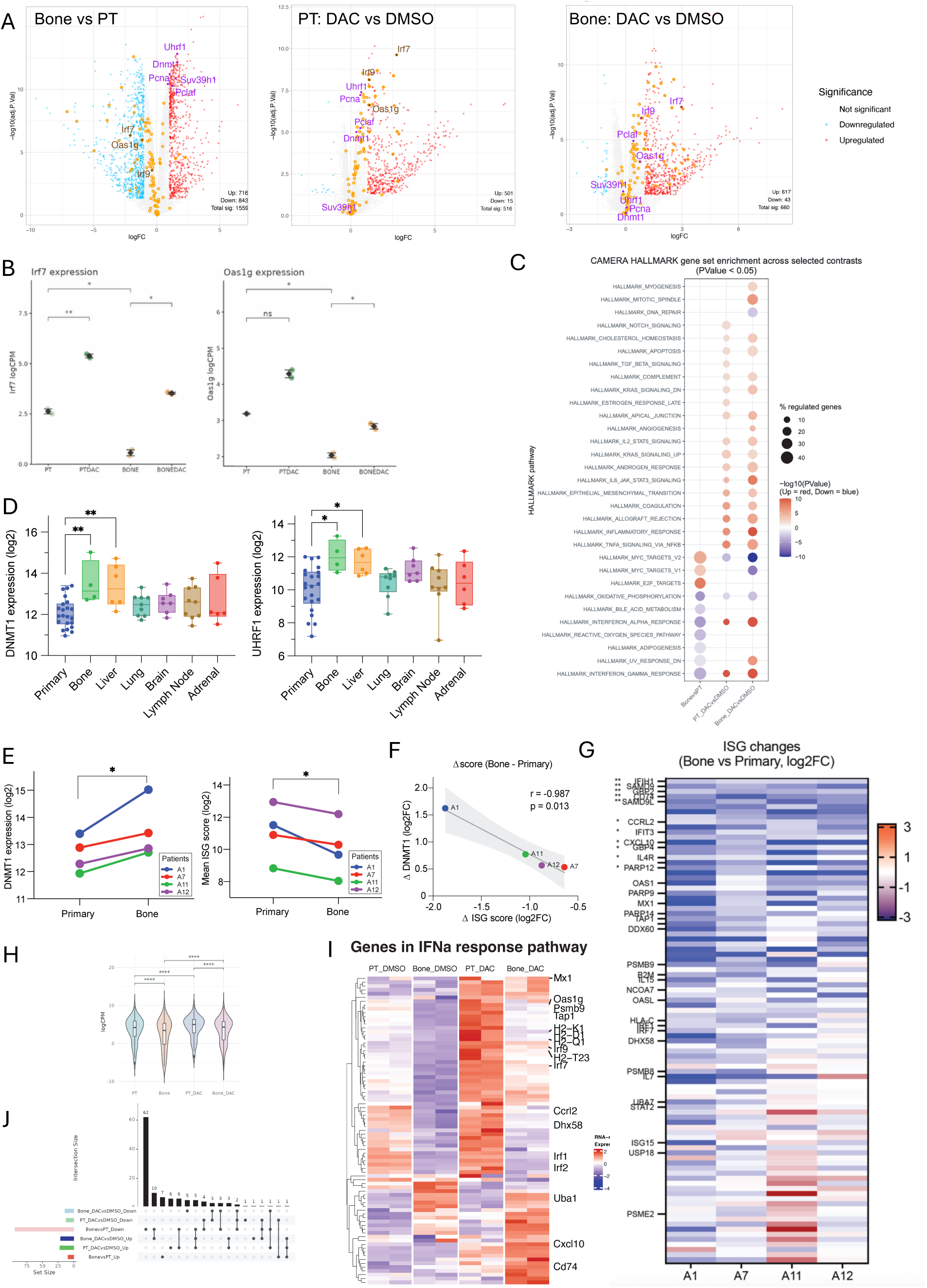
DAC Reactivates Interferon Signalling in Bone- and Primary-Derived Tumor Cells with Distinct Transcriptional Programs. **(A)** Volcano plots comparing Bone and matched PT 4T1.2 cell lines at baseline (left), PT following DAC treatment (middle), and Bone following DAC treatment (right). Yellow circles represent ISGs. Selected ISGs and DNMT1 related genes are highlighted. **(B)** Expression of representative ISGs across PT and Bone cell lines with or without DAC treatment. **(C)** Gene set enrichment analysis (GSEA) comparing Bone and PT cells, with or without DAC treatment. **(D)** Expression of DNMT1 and UHRF1 across primary tumors and metastatic sites in the RAP cohort from the AURORA cohort. Bone metastases exhibited the highest expression of both methylation maintenance factors relative to primary tumors. **(E)** Expression of DNMT1 (left) and ISG score (right) in matched primary tumor and bone metastasis samples from the AURORA cohort. Bone metastases demonstrated increased methylation machinery expression accompanied by reduced ISG expression. **(F)** Correlation analysis between DNMT1 expression and ISG scores in bone metastasis samples. Increased DNMT1 expression was associated with reduced interferon signalling. **(G)** Heatmap showing differential expression analysis of 95 curated ISGs in matched primary tumor and bone metastasis samples. Eighteen ISGs were significantly downregulated in bone metastases, with additional *IRF7*, *IRF1* and IFN-responsive genes including *OAS1*, *MX1* and *TAP1* showing consistent suppression. **(H)** Violin plots showing gene expression in IFNα response pathways following DAC treatment in PT and Bone cells. DAC robustly reactivated interferon signalling in both cellular contexts, although the magnitude of pathway activation was reduced in Bone cells. **(I)** Heatmap showing genes in IFNα pathway activation following DAC treatment. PT cells displayed strong IFNα induction compared with Bone cells, indicating context-dependent responsiveness to DNA hypomethylation. **(J)** Overlap of DAC-responsive genes in PT and Bone cells. Approximately one-third of differentially expressed genes were commonly showing differential expression accross cell types, demonstrating substantial tissue-specific transcriptional responses to DAC treatment. Data represented as mean with min to max for B, D and H, P-values were calculated using two-tailed, unpaired, non-parametric t-test. * P < 0.05, **P < 0.01, *** P < 0.001, **** P < 0.0001. ISG, interferon-stimulated gene; GSEA, gene set enrichment analysis.

### Clinical validation of methylation-associated transcriptional reprogramming in bone metastasis

To determine the clinical relevance of these findings, we interrogated BCa transcriptomic data from the RAP cohort of the AURORA US network that utilised rapid autopsy samples from a range of tissues (GSE193103). Across metastatic sites, bone metastases exhibited the highest expression of both *DNMT1* (>2.7-fold increase, p = 0.006) and *UHRF1* (∼4-fold increase, p = 0.006) relative to primary tumors (Fig. 3D).

Analysis restricted to paired patient primary-bone metastases samples confirmed consistent upregulation of *DNMT1* and *UHRF1* in bone lesions, accompanied by suppression of a subset of ISGs (Fig. 3E). Although statistical power was limited by sample size, bone metastases demonstrated the lowest overall ISG expression trend compared with other metastatic sites, and highly correlated with the expression of *DNMT1* (Fig.3F). Notably, 18 of 95 analysed ISGs were significantly downregulated, while additional key ISGs including *OAS1, MX1,* and *TAP1* showed consistent suppression across matched samples (Fig. 3G), supporting the clinical relevance of our 4T1.2 subline transcriptional data. Collectively, the reciprocal pattern of suppressed IFN programme with increased DNMT1 maintenance methylation machinery in bone suggests DNA methylation may contribute to ISG suppression as an immune evasion mechanism in bone metastases.

### DAC reactivates interferon programmes but reveals site-specific transcriptional responses

To investigate whether DNA methylation contributes to IFN pathway suppression and whether the DNMT inhibitor DAC can rescue this loss of ISGs, Bone and PT cell lines were treated with the hypomethylating agent DAC. Differential gene analysis confirmed robust reactivation of interferon signalling pathways following DAC treatment in both PT and bone cells (Fig. 3A). Key interferon regulators and ISGs, including *Irf7*, were significantly induced following DAC treatment in both cell contexts (Fig. 3A-B).

Interestingly, DAC treatment induced distinct interferon effector programmes in PT versus Bone cells. While PT cells displayed stronger global IFNα pathway activation, Bone cells demonstrated a quantitatively delayed response (Fig. 3H-I). In fact, only approximately one-third of DAC-responsive genes overlapped between Bone and PT contexts (Fig. 3J), suggesting that tissue-specific epigenetic wiring influences IFN pathway reactivation.

Together, these findings indicate that DAC can restore particular type I IFN signalling programs in bone-derived tumor cells, the magnitude and composition of which is tissue source dependent. This suggests that DAC may act through alternative mechanisms in different contexts. We therefore next want to confirm whether this bone-specific effect could be attributed to altered DNA methylation at these genes across the different cell types.

### Divergent baseline methylation landscapes between Bone and PT tumor cells

To define the epigenetic basis of transcriptional divergence, we profiled genome-wide DNA methylation using Infinium Methylation EPIC array across all cell lines with and without DAC treatment. Principal component analysis demonstrated clear separation between Bone, PT, and Parental methylation profiles, confirming stable site-specific epigenetic reprogramming. Notably, PT cells remained epigenetically distinct from the parental cell line, suggesting persistent tissue-derived methylation states despite prolonged *in vitro* culture.

At baseline, comparison between Bone and PT cells identified widespread differential methylation (Fig. 4A), with 3.58% of probes hypomethylated and 6.67% hypermethylated in Bone cells (Fig. 4B-C). Differentially methylated probes were modestly enriched within intronic regions compared with promoters, exons, UTRs, and CpG island-associated regions (Fig. 4C).

**Figure 4.**
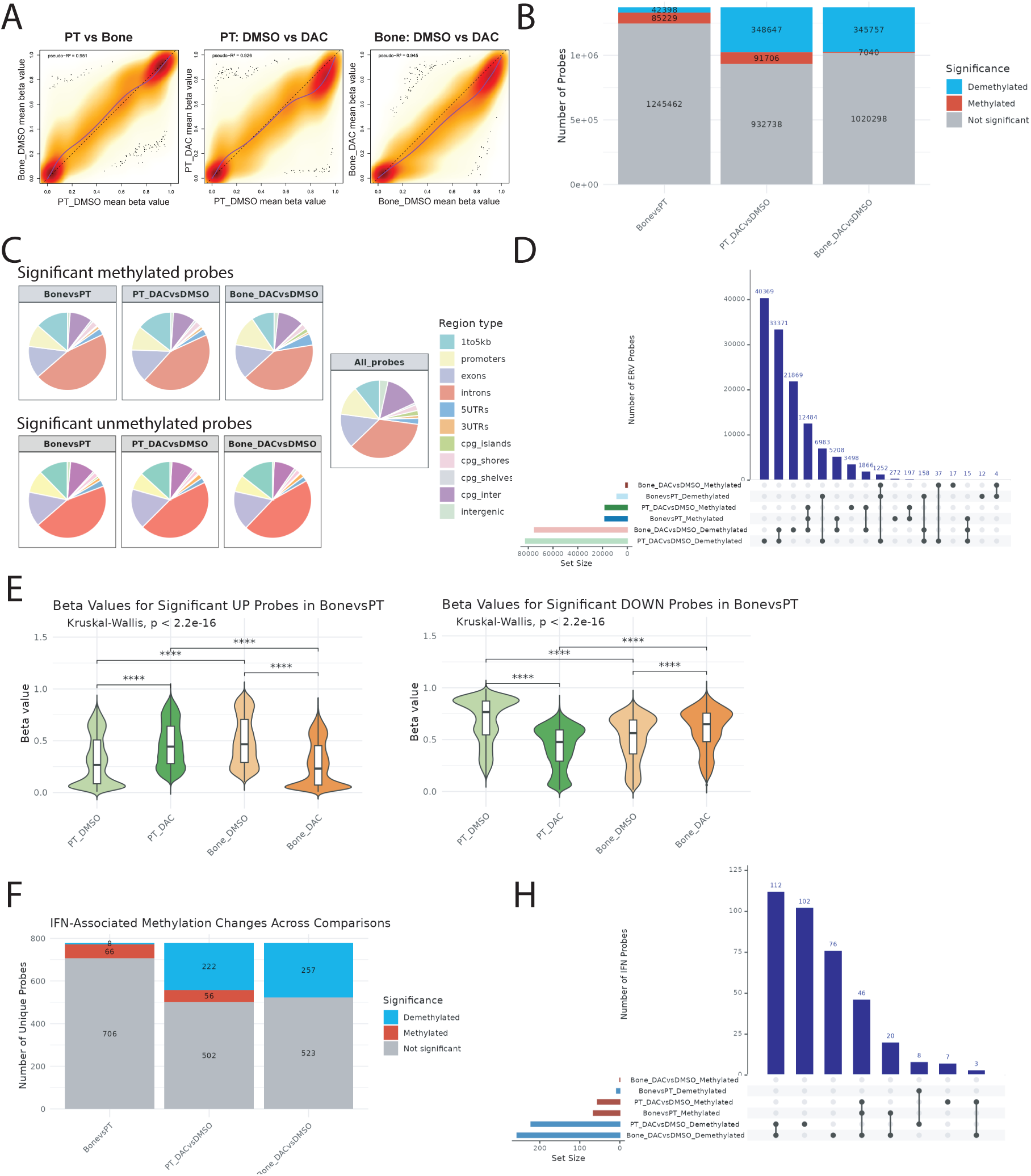
Bone and PT Cells Exhibit Distinct and Plastic DNA Methylation Landscapes Revealed by DAC Treatment. **(A)** Correlation heatmap showing global DNA methylation profile readouts from EPIC arrays in PT and Bone (left), following vehicle or DAC treatment in PT (middle) and Bone (right). **(B)** Distribution of differentially methylated probes identified between PT and Bone cells at baseline and following DAC treatment. **(C)** Genomic annotation of differentially methylated probes identified between different comparisons. Differential methylation was observed across multiple genomic features, with modest enrichment within intronic regions relative to promoters, exons, untranslated regions, and CpG island-associated regions. **(D)** Overlap of DAC-responsive methylation changes among PT and Bone cell lines. While a subset of DAC-responsive probes was shared, substantial cell type-specific methylation remodelling was observed, particularly for the demethylation in PT or in Bone cells. **(E)** Violin plots showing relationship between baseline differential methylation and DAC-induced methylation changes. Probes hypermethylated in Bone relative to PT overlapped with loci gaining methylation following DAC treatment in PT cells (left), whereas probes hypomethylated in Bone frequently corresponded to loci gaining methylation following DAC treatment in Bone cells (right). **(F)** Distribution of differentially methylated probes at ISG-associated loci. Bone cells exhibited enrichment of hypermethylated ISG-associated probes relative to PT cells at baseline. Following DAC treatment, PT cells displayed both methylation gain and loss across ISG loci, whereas Bone cells predominantly underwent demethylation. **(G)** Overlap of ISG-associated probes differentially methylated between Bone and PT cells and probes altered following DAC treatment. The majority of ISG-associated probes hypermethylated in Bone cells overlapped with loci gaining methylation in PC cells following DAC treatment. Data represent biological replicates as described in Methods. Differentially methylated probes were identified using the statistical thresholds specified in Methods. Data represented as mean with min to max for E and I, P-values was calculated using two-tailed, unpaired, non-parametric t-test. * P < 0.05, **P < 0.01, *** P < 0.001, **** P < 0.0001.

### DAC induces global demethylation but reveals context-dependent methylation plasticity

DAC treatment induced extensive genome-wide methylation loss across all cell lines, affecting approximately 27.9 - 32.5% of probes (Fig. 4A-B). Interestingly, PT cells uniquely exhibited a subset of probes demonstrating increased methylation following treatment (6.72%). Although partial overlap existed among DAC-responsive probes across cell lines, a substantial fraction of methylation changes remained cell-type-specific, particularly in Bone cells, where 23.2% of demethylated probes were uniquely altered following DAC exposure (Fig. 4D).

Remarkably, 70.3% of genomic regions that gained methylation following DAC treatment in PT cells corresponded to regions already hypermethylated in untreated Bone cells relative to PT (69.8% of all Bone-hypermethylated regions) (Fig. 4D-E). Conversely, DAC treatment induced a hypermethylation profile in PT cells that substantially overlapped with the hypermethylation signature present in untreated bone metastatic cells. Although the mechanistic implications of these hypermethylated genes have not been tested, this overlap suggests that context and timing are important considerations when applying DNMT inhibitors, as DAC exposure may shift primary tumor cells toward a methylation state resembling that of metastatic cells at a subset of loci.

These findings indicate that specific genomic regions exhibit pronounced methylation plasticity, capable of both methylation gain and loss depending on cellular context and pharmacologic perturbation.

### ISG-associated loci exhibit context-specific methylation regulation

Analysis restricted to ISG-associated probes revealed enrichment of methylation gain in Bone cells relative to PT (8.46 %; Fig. 4F). Following DAC treatment, PT cells displayed both methylation gain (7.18%) and loss (28.46%) across ISG loci, whereas Bone cells exhibited predominantly demethylation (32.9%) (Fig. 4F). Consistent with genome-wide observations, the majority of hypermethylated probes in Bone relative to PT overlapped with loci gaining methylation following DAC treatment in PT cells (66.7% vs 82.1%; Fig. 4H).

### Integration of DNA Methylation and Transcriptomic Profiles Reveals Context-Dependent Epigenetic Control of IFN Signalling

To investigate how DNA methylation contributes to IFN pathway regulation, we integrated EPIC DNA methylation and RNA-seq datasets across PT and Bone cell lines, with and without DAC treatment. We first assessed the global relationship between gene expression and DNA methylation by stratifying methylation probes according to genomic annotation. Across all expressed genes, DNA methylation levels showed weak or no correlation with transcript abundance, regardless of genomic regions (Fig. 5A), indicating that methylation-expression coupling is not a dominant global feature in these cells. In contrast, when the analysis was restricted to ISGs (n=128), promoter-proximal methylation (Promoters; within 1 kb of transcription start sites) exhibited the strongest inverse correlation with gene expression (Fig. 5B), suggesting that promoter methylation plays a selective regulatory role in IFN signalling.

**Figure 5.**
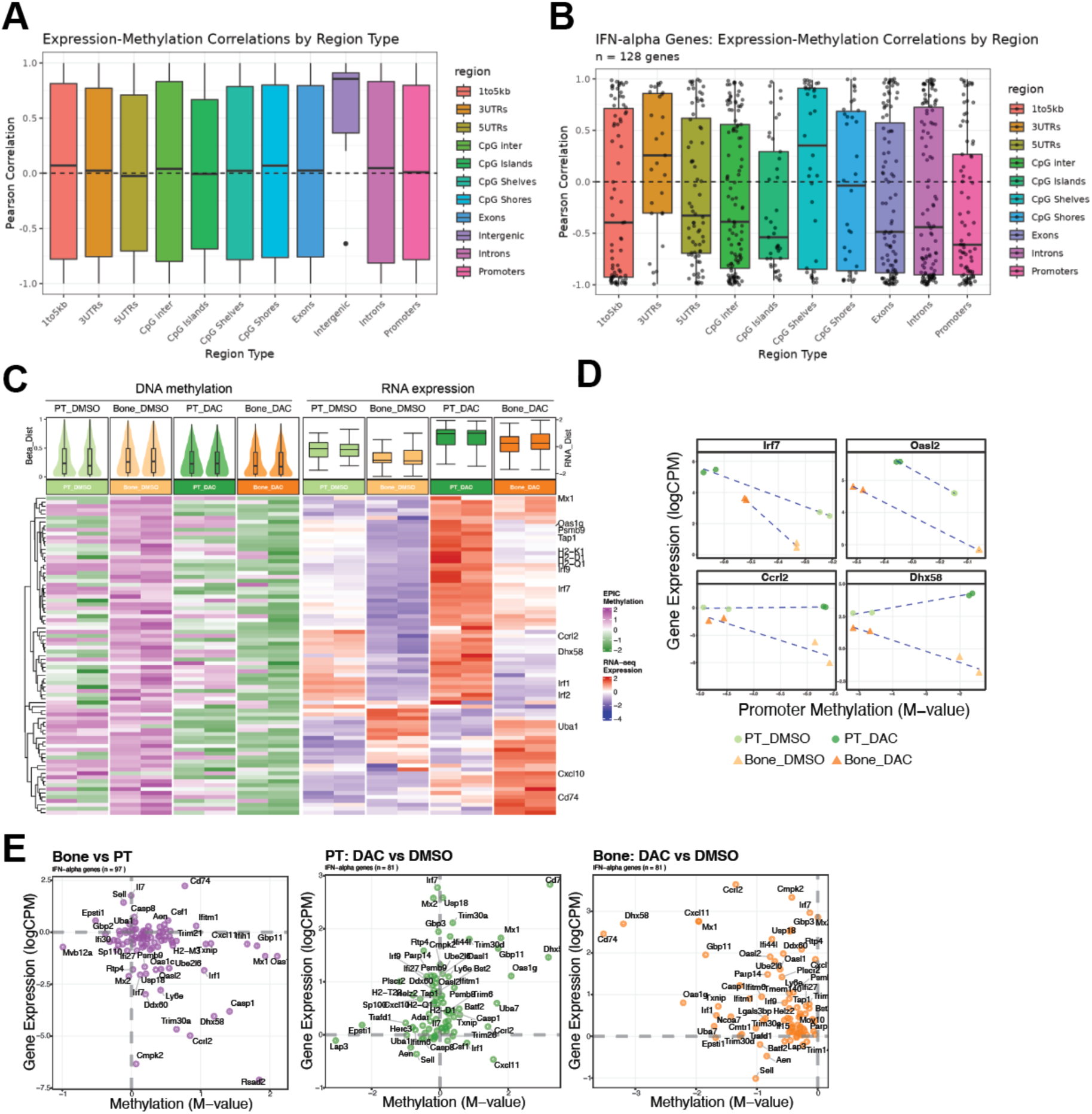
DNA Methylation and Transcriptomic Profiles Reveals Context-Dependent Epigenetic Control of IFN Signalling. **(A)** Correlation between gene expression and DNA methylation at different genomic regions, showing weak or no correlation with transcript abundance, regardless of genomic regions. Result for the Intergenic region is not conclusive as only 5 probes are located within this region. **(B)** Correlation of the ISG genes’ expression to the level of methylation at different genomic regions. Promoters (within 1 kb of transcription start sites) exhibited the strongest inverse correlation with gene expression. **(C)** Heatmap of the DNA methylation at the promoter regions (left) and transcriptomic Profiles (right) of ISGs in PT and Bone cells upon DAC exposure. The heatmap on the RNA expression is identical to Figure 3I. The two columns of each group represent the biological replicate samples. **(D)** The level of promoter methylation and gene expression of representative ISGs across PT and Bone cell lines with or without DAC treatment. **(E)** Correlation plot of differential DNA methylation and gene expression of ISGs in Bone relative to PT (left), PT following DAC treatment (middle) and Bone cells following DAC treatment (right).

Based on this observation, we focused subsequent analyses on promoter-associated methylation of ISGs. ISGs that were transcriptionally suppressed in bone-derived cells at baseline, including *Irf7* and *Oasl2*, showed increased promoter methylation relative to PT-derived cells, consistent with methylation-mediated repression (Fig. 5C). In contrast, ISGs with elevated expression in bone did not display corresponding promoter methylation changes, indicating that additional regulatory mechanisms contribute to their activation.

Following DAC treatment, the majority of ISGs re-expressed in both PT- and bone-derived cells exhibited reduced promoter methylation, supporting a functional link between demethylation and transcriptional reactivation (Fig. 5C-D). Notably, a subset of ISGs that were suppressed in bone at baseline, such as *Ccrl2* and *Dhx58*, became transcriptionally induced in both PT and bone contexts despite gaining promoter methylation in PT cells upon DAC exposure. Similarly, ISGs that were selectively induced in bone following DAC treatment often exhibited increased promoter methylation in PT cells, underscoring a context-dependent and non-linear relationship between promoter-specific methylation and gene expression.

Importantly, changes in ISG expression correlated with promoter demethylation specifically in bone-derived cells, but not in PT-derived cells, indicating that DNA methylation contributes to IFN pathway regulation preferentially in epigenetically reprogrammed metastatic contexts rather than in the original tumor lineage (Fig. 5E).

### Post-resection DAC treatment reduces metastatic spread of 4T1.2 in vivo

Our previous data suggest that treatment of bone metastases with DAC could reverse IFN silencing as a promising anti-metastatic therapeutic. To investigate the *in vivo* efficacy of DAC as an anti-metastatic therapy in an aggressive immunocompetent model, mice were injected with 4T1.2 cells into the mammary fat pad and tumors resected after 14 days, before randomisation into DAC or DMSO groups. Treatment was administered via intraperitoneal injection three times on days 18, 20 and 22 before endpoint on day 23 (Fig. 6A). Bioluminescence imaging (BLI) at endpoint (Fig. 6B) revealed dramatic reductions in metastasis in the DAC treated groups, including an 85% reduction in lung (Fig. 6C) and spine metastasis (Fig. 6D). Flow cytometry analysis of peripheral blood collected from the end point revealed a reduction in the total number of immune cells in the DAC treated group (Fig. 6E). Furthermore, analysis of ratios of lymphoid to myeloid cells in the blood revealed a switch from 25.8:74.2 in the DMSO treated group to 79.1:20.9 in the DAC group (Fig. 6F) outlining a collapse of the myeloid compartment with therapy. Within the myeloid cell ratios, there was an increase in the percentage of basophils and decrease in Ly6G+ neutrophils, the latter aligning with a MDSC cell population previously documented to facilitate metastasis in this model. Together, these data support DAC as a potent anti-metastatic therapeutic in vivo.

**Figure 6.**
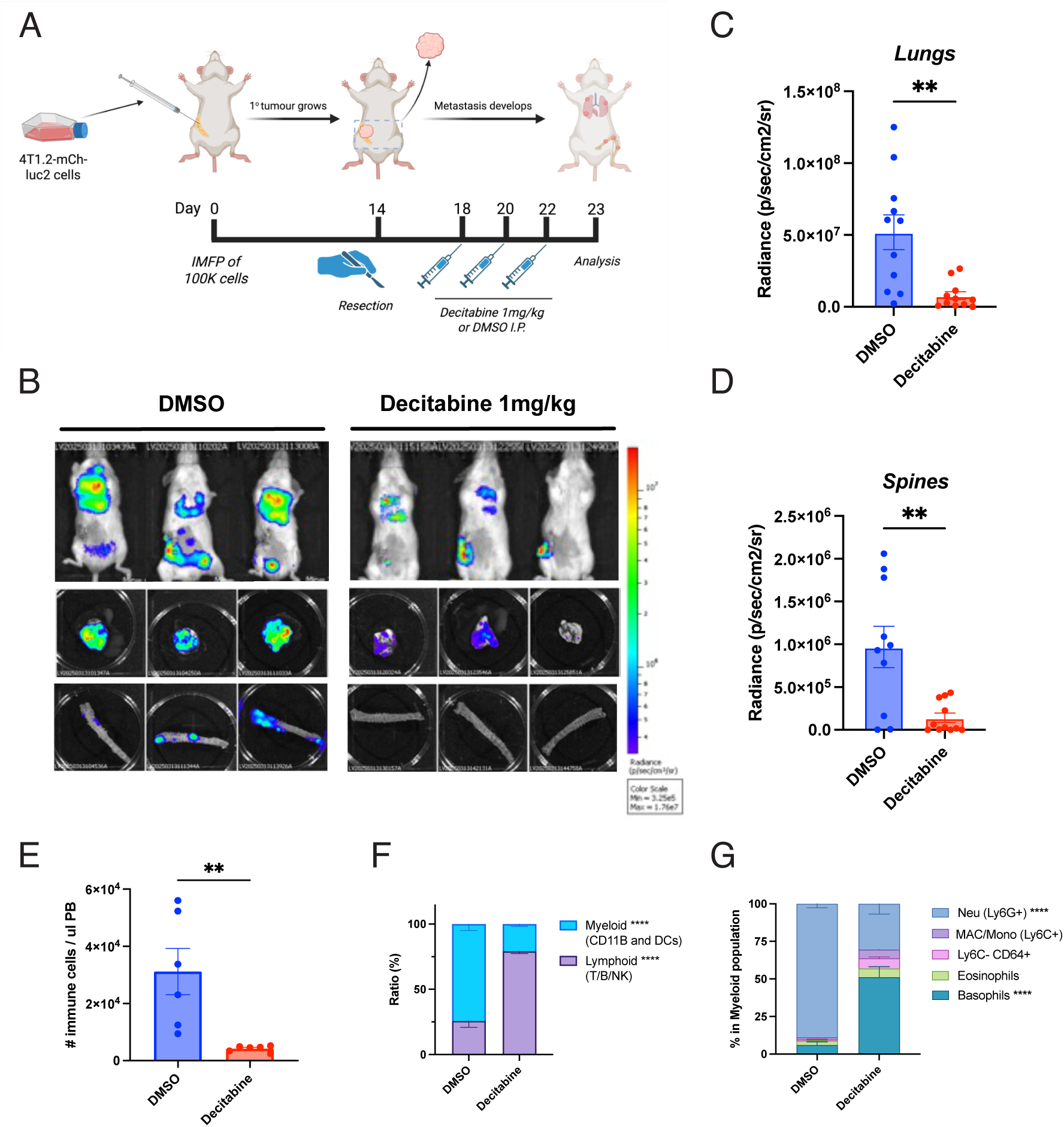
In vivo adjuvant DAC treatment reduces metastasis in the 4T1.2 model and results in myelosuppression. **(A)** Experimental workflow of Balb/c mice treated with DAC or DMSO post 4T1.2 intramammary fat pad (IMFP) resection to simulate adjuvant therapy *in vivo*. **(B-D)** Bioluminescent imaging to quantify 4T1.2 tumor burden in lung **(C)** and spine **(D)**. **(E)** Flow cytometry of the peripheral blood of treated mice analysed total immune cells in the DAC group. **(F)** The ratios between lymphoid (T(TCRb+), B(CD19+B220+) and natural killer (NK; CD49b+)) and myeloid (CD11b^+^ and dendritic cells (CD11c+, DCs)). **(G)** The ratios of Ly6G+ neutrophils, Ly6C+ macrophages/monocytes, Ly6C- CD64+ cells, eosinophils and basophils. Data represented as mean +/- SEM. Statistical a two-tailed, unpaired, nonparametric t-test. N =10-11 for **(C, D)** and n = 6 for **(E-G).** P-values was calculated using two-tailed, unpaired, non-parametric t-test. * P < 0.05, **P < 0.01, ****P < 0.0001.

## DISCUSSION

Once bone metastasis becomes clinically evident, it is largely incurable^41^. Substantial evidence underscores the importance of reactivating IFN signalling in bone-metastatic cancer, both to enhance response to chemotherapy and to restore immunogenicity for re-sensitization to immunotherapy. Here, we identify several clinically available compounds, including DAC, Mocetinostat, and DZNep-HCl, that consistently induce ISRE reporter activity in a dose-dependent manner, with DAC the only compound that does not affect cell viability, indicating its potential for boosting the immunogenicity of viable cells (Fig. 1A-B). Notably, we also observed that 4T1.2_BONE cells exhibit increased sensitivity to the Aurora kinase inhibitors Barasertib and SNS-314 mesylate compared with matched 4T1.2_PT cells, indicating a bone-specific response. This is of particular interest given that we observed MYC amplification in BONE cells (Fig. 3C), and Aurora kinase acts downstream of MYC signalling, raising the possibility of targeting Aurora kinase to boost immunogenicity. However, because DAC confers broad induction of ISRE signalling across a wide range of cell lines, we focused our subsequent investigation on DAC.

DNMT mediated repression of ISGs is previously well described in models of BCa (as described in the introduction). This has important ramifications for resistance to newer therapies such as immune checkpoint blockade (ICB), where it has been described that DNMT-mediated methylation of STING, IRF7 and PD-L1 may confer resistance to therapy^31,42^. Indeed, the discovery of DNMT inhibition and the stimulation of IFN signalling through transcription of endogenous retroviral elements (ERVs), short interspersed nuclear elements (SINEs), long interspersed nuclear elements (LINEs) and other retroelements has opened the avenue of DNMT inhibitors like DAC as a IFN inducer as well as a de-repressor of gene methylation^38,39,43–45^. There has therefore been extensive use of DAC in preclinical BCa models to as a single agent^46–48^, in combination with doxorubicin^49^, with ICB^42,50^ and even anti-cancer vaccines^51^ (reviewed in ^52^). DAC has even shown promise in the clinic in multiple clinical trials, most recently the phase 2 study (NCT02957968) which reported a pathological complete response in 11/27 (40.1%) when patients were treated with DAC and pembrolizumab (anti-PD-1) before neoadjuvant chemotherapy^53^. This was confirmed in our results, as all parental and primary tumor cell lines demonstrated IFN pathway induction in response to DAC alone.

While DAC has shown anti-metastatic potential in 4T1 syngeneic^46,54,55^ and MDA-MB-231 xenograft models in vivo^55,56^, its use as an IFN-sensitising therapy for bone metastasis had previously been unexplored. This is relevant not only due to the frequency and treatment resistance of bone metastasis in BCa patients but also due to our previous discovery of widespread IFN and ISG downregulation in bone metastatic lesions^16^ (further confirmed in Figures 2B). Furthermore, the use of DAC in an anti-metastatic therapeutic context targeting the mechanistic process of type I IFN pathway downregulation may avoid chronic inflammation and therapeutic toxicities. Chronic upregulation of pSTAT, cGAS-STING and ISG signatures have been associated with poorer outcomes in BCa^57–60^ and this has contributed to the limited success of IFN-based therapies over the last 50 years (as reviewed in ^14^). In patients, DAC may sensitise bone metastatic lesions to traditional treatments such as chemotherapy or radiotherapy that induce an IFN response with reduced toxicities compared to the primary setting^14,61^. However, our data showed DAC alone was able to significantly reduce lung and bone metastasis when mice were treated post-resection, outlining its potential as both an IFN-sensitizer and inducer due to the production of dsRNA.

Beyond therapeutics, this study characterised, for the first time, the expression and DNA methylation landscape in bone metastasis compared to the primary tumor. Our laboratory has previously characterized loss of IRF7 and IRF9, and the associated suppression of ISG signatures, in bone metastasis of breast and prostate cancer in both mouse models and patient-derived samples^16–19^. Our RNA sequencing data recapitulate this phenotype and extend it to a broader ISG panel, including the reduction of *Irf1*, a transcription factor that facilitates opening of ISG loci for transcription^62^, and the elevation of *Uba1*, a negative regulator of interferon responses via degradation of JAK1^63^, both of which further restrict IFN signalling in Bone relative to PT (Fig. 3I).

Beyond confirming ISG suppression, we identify a reciprocal upregulation of *Dnmt1* and its associated maintenance methylation machinery (*Uhrf1, Pcna* and *Pclaf*) specifically in bone-derived cells. Elevated DNMT1 has previously been linked to metastatic potential in BCa broadly^64,65^, but prior work on epigenetic immune evasion has largely been focused on soft-tissue metastasis^66,67^.

Given that bone is the most common site of BCa metastasis, the absence of site- specific mechanistic data represents a notable gap. To our knowledge, this is the first demonstration of a reciprocal DNMT1-ISG anti-correlation specifically in bone-metastatic cells relative to matched PT, in both a syngeneic mouse model and patient samples. Cha *et al.* reported that DNMT1 was downregulated in bone metastasis overall, however, this phenotype is subtype-dependent, with DNMT1 upregulated in basal and HER2+ subtypes, where its elevation was associated with significantly worse survival^63^. Supporting the clinical relevance of our findings, paired primary-bone metastasis samples from the RAP cohort showed the same pattern, with elevated DNMT1 strongly correlated with suppressed ISG expression in bone (Fig. 3D-G), indicating this is not a mouse-model artefact but a feature also present in patients. Whether this phenotype is driven by cellular crosstalk within the bone marrow microenvironment, or is cell-intrinsic to bone-tropic tumor clones, remains to be determined at this resolution.

Relative to PT, loss of ISG expression in bone is strongly correlated with gain of methylation at gene promoter regions with reduced expression magnitude (Fig. 5C). This nonlinear relationship between methylation loss and reactivation of individual genes indicates that additional partner factors constrain the magnitude of ISG induction in the Bone cells. For instance, *Uba1*, a negative regulator of ISG, is more highly expressed in the Bone, where it likely desensitises the signalling cascade by limiting availability of JAK1, independent of the loss of methylation at the Uba1 promoter itself. This persistence of negative regulators such as Uba1 may partly explain why ISG suppression in bone cannot be fully reversed by demethylation alone. Whether this reduced magnitude of ISG expression is sufficient to open a therapeutic window for bone-metastatic lesions, or whether it limits DAC’s utility as a monotherapy, has not previously been addressed.

Our analysis focused specifically on promoter methylation; we did not examine other genomic regions (for instance: CpG islands, gene bodies), which may offer a complementary explanation for this phenotype (Fig. 5B). Promoter methylation governs expression at a majority of ISG loci, but regulatory methylation at other genomic elements could independently constrain expression at the remaining loci, such that even where DAC successfully erases promoter methylation, these genes would not be expected to reactivate. DAC’s incomplete restoration of ISG expression in bone is therefore mechanistically consistent with this model, rather than reflecting a limitation of the drug itself.

Having established that ISG reactivation in bone is attenuated relative to PT, we next asked whether this re-activation with reduced magnitude is nonetheless sufficient to confer a therapeutic benefit *in vivo*. Using the IMFP model, in which DAC treatment was initiated after surgical resection of the established primary tumor, we observed a comparable benefit, indicating that DAC acts directly to limit the progression of disseminated cells in both the spines and in the lungs (Fig. 6B-D). Consistent with other reports^47,68^, we also observed that DAC depletes myeloid-derived suppressor cells in our model, indicating an additional effect on the immune compartment. To delineate tumor-intrinsic from direct immune cell targeting, pre-treating 4T1.2 cells with DAC prior to engraftment is one avenue to explore to clarify whether tumor-intrinsic effects of DAC alone are sufficient to inhibit metastatic progression.

Together, our findings report that metastases derive methylation profiles independent to that of the primary tumor and that this drives loss of key pathways associated with bone metastatic progression. Decitabine is one avenue to rescue the epigenetic loss of type I IFN signalling that could have major metastasis-specific impacts on therapeutic considerations in the future where the aim is to target the cellular plasticity that allowed metastatic outgrowth in new microenvironments.

## METHODS

### Cell culture and reagents

The 4T1.2 mouse BCa cell line is a highly metastatic derivative of the 4T1 line^69^, originally derived from a spontaneous mammary tumor in a female Balb/c mouse^70^. Cells from a matched primary tumor and bone metastatic lesion of the same Balb/c mouse were isolated in house to generate the 4T1.2_PT and 4T1.2_BONE lines. The J110 line is a syngeneic metastatic cell line originally characterised as ER-positive but later confirmed as stromal (ER-negative and tamoxifen-insensitive) derived from the mammary tumor of an MMTV-Amplified in Breast 1 (AIB1)-transgenic mouse^71–73^. The MDA-MB-231, CAL-120 and T47D human cell lines were derived from the pleural effusions of female patients with metastatic breast carcinoma^74–76^.

The cell lines and their fluorescent reporter derivatives were maintained in the following conditions: 4T1.2-derived murine mammary tumor cell lines, were maintained in α Minimal Essential Medium (αMEM; Gibco #12561056) and 5% Foetal Calf Serum (FCS; Hyclone #SH30406); J110 were maintained in Dubecco’s Modified Eagle Medium (DMEM)/Ham’s F12 (Gibco #12500062) and 10% FCS; MDA-MB-231 and CAL-120 were maintained in DMEM, high glucose (Gibco, #11965092) with 10% FCS and 2mM GlutaMAX (Gibco #35050061); T47D were maintained in Rosewell Park Memorial Institute (RPMI)-1640 (Gibco #11875093) with 10% FCS. All cells were incubated at 37°C in 5% CO2 and all media contained penicillin (100 U/ml, Merck #P3032) and streptomycin (100 μg/ml, Merck #S9137)

### Lentiviral generation and cell transduction

To generate a stable ISRE-GFP-luc2 reporter cell lines, lentiviral plasmid mixes containing second generation packaging plasmids psPAX2 (Addgene # #12260) and pCMV-VSV-G (Addgene #8454) with the pGreenFire1-ISRE target plasmid (Systems Biosciences #TR016PA-1) in a ratio dependent on target plasmid size, as described in Stewart et al^77^. These plasmid mixes were transfected into human embryonic kidney-293T (HEK293T) cells with polyethyleneimine (3:1 ratio to total plasmid amount; Kyfora Bio #23966) containing serum and antibiotic-free DMEM. Fresh DMEM + 10% FCS media was added to the HEK-293T cells after 24 hours, and 48-72 hours post transfection, the lentiviral-containing media was added to the target along with 8μg/mL Polybrene (Sigma-Aldrich/Merck #H9268). Cells were selected with puromycin (Sigma-Aldrich/Merck #P9620) containing media.

### Epigenetic compound screen

The epigenetic compound library (CA epigenetic library) comprising of 181 compounds was acquired from Compounds Australia. Four BCa cell lines including 4T1.2, J110, CAL120, and T77D were seeded in 384-well plates (Corning Life Sciences #3764) at different densities (Table 1). The following day they were treated with epigenetic compounds for 72 hours at five concentrations: 5 µM, 1µM, 0.2 µM, 0.05 µM, 0.005 µM. The screen for each compound was performed as one replicate. Nineteen hit compounds identified in the initial screen were further validated in a secondary screen across seven BCa cell lines including MDA-MB-231, CAL-120, T47D, 4T1.2 parental, 4T1.2 bone, 4T1.2 PT, and J110. This validation screen was conducted in replicate at 10 concentrations: 5µM, 3µM, 2µM, 1µM, 0.75µM, 0.5µM, 0.2µM, 0.1µM, 0.05µM, 0.0055µM. ISRE activation was assessed via bioluminescence measuring Luc2 activity using Bright-Glo Luciferase Assay System (Promega, #E2610) with Biotek Cytation5. Viability was assessed using CTG viability assay. Luminescent reads and viability were normalized to negative control, DMSO. Drugs with a z-score equal or above 2 were considered significant.

**Table 1:** Cell lines and seeding numbers for epigenetic compound screen.

| Cell lines | Cells/well |
| --- | --- |
| MDA-MB-231 | 750 |
| CAL-120 | 550 |
| T47D | 1000 |
| 4T1.2 PARENTAL | 350 |
| 4T1.2 BONE | 350 |
| 4T1.2 PT | 350 |
| J110 | 125 |

### Quantification of mRNA expression after Decitabine treatment

To determine the effect of Decitabine treatment on 4T1.2_PT and _BONE cells, cells were treated with 250nM Decitabine (DAC, Selleck Chemicals #S1200) or dimethyl sulfoxide control (DMSO, Merck #317275) for 48 hours. Messenger RNA (mRNA) was extracted as per kit instructions from cell pellets utilising the RNeasy Mini Kit (Qiagen #74106) and treated for DNA contamination with the RNase-Free DNase Set (Qiagen # 79256). Complementary DNA (cDNA) was generated from purified mRNA using the SuperScript™ III First-Strand Synthesis System as per supplier’s instructions (Thermo Fisher #18080051).

Quantitative real time polymerase chain reaction (qRT-PCR) was performed to compare mRNA expression of canonical ISGs *Irf7, Irf9, Oas1, Psmb9* and *Tap1* to housekeeping gene *Rps27a*. Each well in a 384-well plate contained 2μL cDNA, 2x Fast SYBR Green dye (Thermo Fisher #4385616) and 0.2μM of a 10μM forward and reverse primer mix (Table 2, synthesised by Integrated DNA Technologies (IDT)). Samples were run in triplicate and each biological replicate data point was from a separate freeze-thaw and plate run.

**Table 2:**
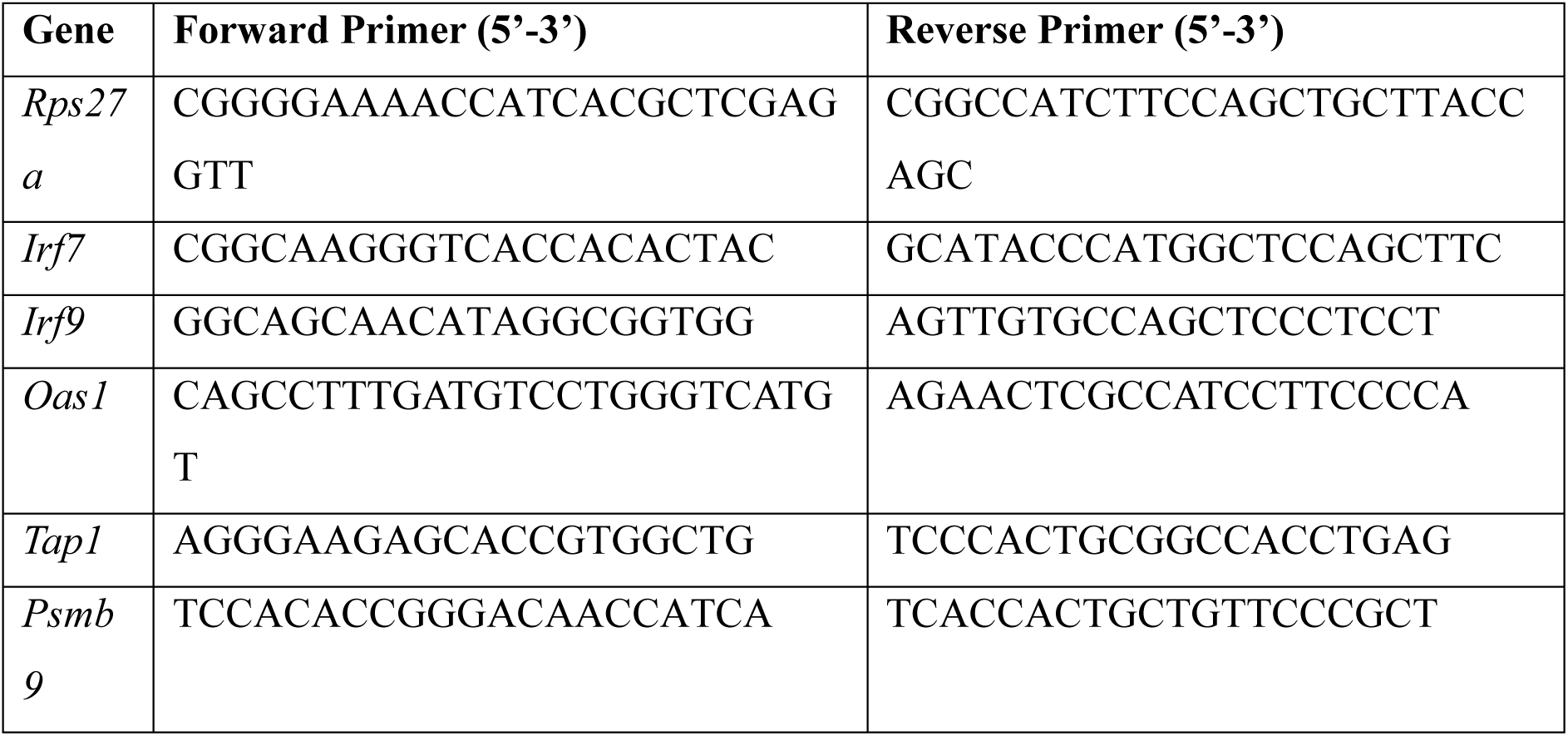
Primer sequences for qRT-PCR.

### Validating Decitabine’s effect on ISRE expression using flow cytometry

4T1.2_PT-ISRE and 4T1.2_BONE-ISRE cells were seeded in 12-well plates and treated with Decitabine for 48-hours and Poly (I:C) (Sigma Aldrich, #P1530) 24-hours prior to harvest. Poly (I:C) was transfected into cells via a 1:3 ratio to Lipofectamine 2000 (Invitrogen #11668027). Cells were harvested and resuspended in FACS buffer containing 2% FCS, 1mM EDTA in phosphate-buffer saline (PBS, Gibco #10010023) and stained with Fixable Viability Dye eFluor™ 660 (1:1000, eBioscience #65-0864-14) to identify live cells. Acquisition was performed on the FACSymphony A5 SE (BD Bioscience) and analysis performed using FlowJo v10.4 (BD Bioscience).

### Flow cytometry analysis of blood

Whole blood (50uL) was incubated in ACK red cell lysis buffer (150 mM NH4Cl, 10 mM KHCO3, 0.1 mM EDTA) for 5 mins at room temperature and washed in FACS buffer (PBS, 2% FBS) twice. Cells were then resuspended in PBS with live-dead stain for 30 mins and washed in FACS buffer twice. Cells were then resuspended in FACS buffer and stained with fluorophore-conjugated antibodies against cell surface markers (detailed list of antibodies is provided in Table 3). Normal mouse bone marrow cells were used as a comparator, and count beads (Beckman Coulter #7547053) were added to each sample after staining to enable quantification of cell numbers. Flow cytometric analysis was performed on Symphony A5 flow cytometers (BD Biosciences), and data were analysed using FlowJo software (Tree Star).

**Table 3:** Antibodies for flow cytometry.

| Cell marker | Final | Company | Cat # | Dilution |
| --- | --- | --- | --- | --- |
| Live dead | FVS700 | BD Horizon | 564997 | (1/1000) |
| FcgR | BUV737 | BD Horizon | 565272 | (1/200) |
| CD45.2 | BUV395 | BioLegend | 564616 | (1/200) |
| TCRb | Pe-Cy5 | BD<br>Pharmingen | 553173 | (1/200) |
| CD19 | PacBlue | BioLegend | 115523 | (1/200) |
| NK1.1 | BUV805 | BioLegend | 108918 | (1/200) |
| Ly6G | BB700 | BD Horizon | 566435 | (1/300) |
| Siglec-F | BV786 | BD Optibuild | 740956 | (1/200) |
| B220 | BUV496 | ThermoFisher | 364-0452-80 | (1/200) |
| CD172a | BV605 | BD Optibuild | 740390 | (1/200) |
| Ly6C | PEcy7 | BD<br>Pharmingen | 560593 | (1/300) |
| F4/80 | APC | BD<br>Pharmingen | 566787 | (1/200) |
| CD11b | BUV661 | BD Horizon | 568345 | (1/200) |
| CD11c | BV480 | BD Horizon | 562949 | (1/200) |
| MHCII | APC-Cy7 | Invitrogen | 47-5321-82 | (1/200) |
| CD103 | PE-CF594 | BD Horizon | 565849 | (1/200) |
| PD-L1 | BV711 | BD Horizon | 568561 | (1/200) |
| CD80 | BUV563 | BD Optibuild | 741272 | (1/200) |
| CD4 | FITC | BioLegend | 100510 | (1/200) |
| CD8 | PE | BD<br>Pharmingen | 553033 | (1/200) |

### Experimental animals

All animal experiments related to the 4T1.2 syngeneic mouse model were performed at the Peter MacCallum Cancer Centre and were approved by the Peter MacCallum Cancer Centre Animal Experimentation Ethics Committee. Wild-type Balb/c and immune deficient NOD.Cg-Prkdcscid Il2rgtm1Wjl/SzJ (referred to as NSG) mice were purchased from the Animal Resources Centre of Peter MacCallum Cancer Centre or the Walter and Eliza Hall Institute of Medical Research (Melbourne, Australia). All experimental mice were housed at the Peter MacCallum Cancer Centre under specific pathogen-free conditions. Animals were group-housed in individually ventilated micro-isolator cages (6 mice per cage) on a 13-hour light / 11-hour dark cycle.

### Transplantation and disease monitoring for the 4T1.2 model

In the case of orthotopic injection to generate murine BCa, 100,000 cells were resuspended in PBS and transplanted into Balb/c recipients by intramammary fat pad injection into the 4^th^ inguinal mammary gland. Animals were closely monitored for clinical signs of disease development (weight loss, lethargy, hunched posture). Primary tumors were resected at 300-400mm^3^ (around day 14 post-engraftment). At the experimental endpoint, mice were euthanised and organs were imaged and collected for histological analysis at a single timepoint. For DAC treatment *in vivo*, mice were treated with 1mg/kg of DAC or DMSO intraperitoneally on days 4, 6 and 8 post tumor resection.

### RNA sequencing and analysis

4T1.2 cells were treated with 125 nM DAC or DMSO for 48 hours. RNA was extracted as previously described. Purified RNA was quantified using the TapeStation 2200 system (Aligent) with RNA HS screentape (Aligent) according to the manufacturer’s instructions, and stored at -80 °C until required. For sequencing, the Peter Mac Molecular Genomics Core used the QuantSeq 3’ mRNA-seq Library Prep Kit for Illumina (Lexogen, Cat#015), according to the manufacturer’s instructions, to generate the sequencing libraries from the purified RNA. 100 bp paired-end reads with a depth of 20 million reads per sample were generated using the Illumina NextSeq500. Sequencing reads were demultiplexed using bcl2fastq (v 2.17.1.14), low quality (Q < 30) reads removed, and trimmed at the 5’ and 3’ ends using cutadapt^78^ (v 1.14) to remove adapter sequences and poly-A-tail derived reads respectively. Sequencing reads were mapped to the mouse reference genome (mm10) using HISAT2^79^ (v 2.1.0) and counted using the featureCounts command of the Subread^80^ (v 1.6.3). Read normalisation and differential gene expression analysis was performed in R (v 4.4.1) using R packages limma^81^ (v 3.60.6) and EdgeR^82^ (v 4.4.2). RCisTarget^83^ (v 1.29.0) was used for motif enrichment analysis. Barcodeplots from limma were used to visualize gene set enrichment and rotating gene set testing was used to test for gene set enrichment. R packages pheatmap (v 1.0.13), ggplot2 (v 4.0.0), ggrepel (v 0.8.1), RColorBrewer (v 1.1-3), ComplexHeatmap (v 2.20.0) and UpSetR (v1.4.0) were used to generate figures.

### Methylation EPIC array and analysis

4T1.2 cells were treated with 125 nM DAC or DMSO for 48 hours. DNA was extracted using the Qiagen DNeasy Blood & Tissue Kit as per manufacturer’s instructions (Qiagen, Cat#69504). Purified DNA was quantified using the Qubit dsDNA HS assay (Invitrogen, Cat#Q32851) according to the manufacturer’s instructions, and stored at -20 °C until required. For methylation profiling, Illumina Infinium Mouse Methylation BeadChip arrays was used according to the manufacturer’s instructions. Arrays were scanned using the Illumina iScan platform and raw IDAT files were imported into R (v4.4.1) for preprocessing and quality control using the package SeSAMe^84^ (v1.14.2). Raw signal intensities were processed using the SeSAMe pipeline with Prep Code TQCDPB, which incorporates background subtraction, dye-bias correction, and probe-level quality assessment. Detection performance was evaluated using SeSAMe detection metrics. Quality control metrics, including methylated and unmethylated signal distributions, probe detection rates, and sample clustering, were visualized using ggplot2, ggpubr, patchwork.

To identify differentially methylated regions, CpG statistics were aggregated using the SeSAMe DMR function to identify spatial regions of differential methylation by grouping adjacent CpG sites with consistent directionality of effect. Separate differential analyses were performed for each biological comparison, including Bone versus PT, Bone DAC versus DMSO and PT DAC versus PT DMSO conditions. All results were stored as SeSAMe output objects for downstream interpretation and visualization. Differential methylation analyses were also conducted using the limma package in separate, with statistical significance adjusted for multiple testing using the Benjamini-Hochberg false discovery rate (FDR) method. R package ggplot2 (v4.0.0), ggpubr (v0.6.0), ggthemes (v5.1.0), dplyr (v1.1.4), tidyr (v1.3.1), reshape (v0.8.10), tibble (v3.2.1), forcats (v1.0.0) and UpSetR (v1.4.0) for structured handling of methylation matrices and sample metadata, and generate figures. Colour schemes were applied using viridis (v0.6.5) and RColorBrewer (v1.1-3) to ensure perceptually uniform and publication-ready visualisation. Genomic interval-based visualisation of CpG loci and differentially methylated regions (DMRs) was supported by GenomicRanges (v1.56.2) from the Bioconductor framework.

## DATA AVAILABILITY

Processed and unprocessed data for RNAseq and Methylation EPIC array will be deposited to GEO and accession numbers provided prior to publication. All other data is available from the corresponding author on request.

## ACKNOWLEDGEMENTS

We thank members of the Victorian Centre for Functional Genomics (Jennii Luu and Robert Vary), animal, molecular genomics, and flow cytometry core facilities at the Peter MacCallum Cancer Centre for technical support. This work was supported by research grants to BSP from the National Breast Cancer Foundation (IRES-23-021, 2025/RPG0108) and the National Health and Medical Research Council (Investigator ID 2018167 and Ideas ID 2012943).

The Victorian Centre for Functional Genomics (KJS: RRID:SCR_025582) is funded by the Australian Cancer Research Foundation (ACRF), Phenomics Australia (https://ror.org/0201hm243), through funding from the Australian Government’s National Collaborative Research Infrastructure Strategy (NCRIS) program, the Peter MacCallum Cancer Centre Foundation and the University of Melbourne Collaborative Research Infrastructure Program. We acknowledge Compounds Australia (RRID:SCR_024565) for their provision of specialized compound management and logistics services.

## AUTHOR CONTRIBUTIONS

Conceptualization and design: MF, JS, TBC and BSP. Experiments and data analysis: JS, TBC, MF, LHC, HC, AT, NH, BM, AI, KJC, and BSP. Technical expertise and essential reagents: BM, RP, CS, KJC, KJS. Supervision: JS, TBC, KJS and BSP. Writing and editing: JS, TBC and BSP.

## DECLARATION OF INTERESTS

BSP is Chief Scientific Officer and a shareholder of AlleSense (Australia). This position has no influence on this publication.

