## Supplemental Figure 1 for "Targeting DNA Methylation Reactivates Type I Interferon Signalling in Bone-Metastatic Breast Cancer"

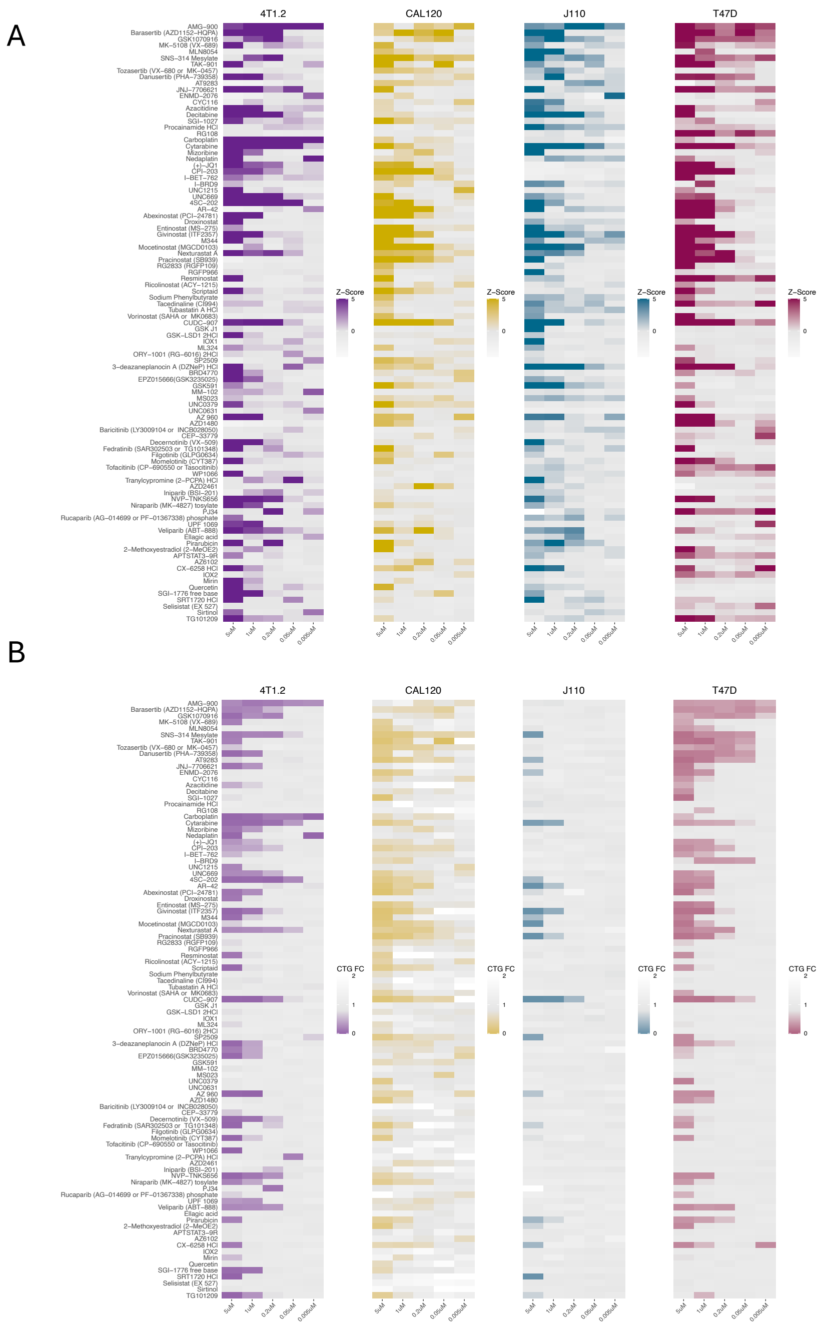


**Supplementary Figure 1. Pharmacological screening of 181 compounds identifies context-dependent inducers of IFN signalling in BCa models.**

**(A)** Heatmap showing ISRE reporter activity following treatment with the indicated compounds for 72 hours. Values represent fold-change relative to vehicle-treated controls.

**(B)** Corresponding heatmap showing cell viability following treatment with the indicated compounds for 72 hours. Values are expressed relative to vehicle-treated controls.
